# The effectiveness of Large Language Models with RAG for auto-annotating phenotype descriptions

**DOI:** 10.1101/2024.11.24.625102

**Authors:** David Kainer

## Abstract

Ontologies are highly prevalent in biology and medicine and are always evolving. Annotating biological text, such as observed phenotype descriptions, with ontology terms is a challenging and tedious task. The process of annotation requires a contextual understanding of the input text and of the ontological terms available. While text-mining tools are available to assist they are largely based on directly matching words and phrases and so lack understanding of the meaning of the query item and of the ontology term labels. Large Language Models (LLMs), however, excel at tasks that require semantic understanding of input text and therefore may provide an improvement for the auto-annotation of text with ontological terms. Here we describe a series of workflows incorporating OpenAI GPT’s capabilities to annotate *Arabidopsis thaliana* and forest tree phenotypic observations with ontology terms, aiming for results that resemble manually curated annotations. These workflows make use of an LLM to intelligently parse phenotypes into short concepts, followed by finding appropriate ontology terms via embedding vector similarity or via Retrieval-Augmented Generation (RAG). The RAG model is a state-of-the-art approach that augments conversational prompts to the LLM with context-specific data to empower it beyond its pre-trained parameter space. We show that the RAG produces the most accurate automated annotations that are often highly similar or identical to expert-curated annotations.

**Short description:** Large Language Models excel at tasks that require semantic understanding of text. Here we use that capability to auto-annotate plant phenotypes with ontological terms and compare to expert annotation.

## Introduction

Large Language Models (LLMs) such as GPT, Claude, LLaMa and Gemini hold great promise for solving problems in many fields. However, caution has been advised when using LLMs for fact-based tasks such as scientific writing. This is due to their generative nature, which can lead to “hallucinatory” effects where the model yields plausible sounding, yet incorrect outputs [1]. Nevertheless, there are certain tasks where LLMs excel and may provide solutions of a quality well beyond the capabilities of other models or algorithms. One such task is the annotation of text with ontological terms [2, 3].

Ontologies are highly prevalent in biology and medicine. The NCBO BioPortal, for example, maintains a library of over 1000 biomedical ontologies. Ontologies are useful for uniting inconsistent information from wide sources under a common lexicon. Once data is annotated with ontological terms, it can be used for classification, simplification/summarizing, data aggregation and over-representation analysis (enrichment) [4, 5]. The process of annotation, however, requires a contextual understanding of the input text and of the ontological terms available. For domain experts the task can be daunting when there is a large body of items to be annotated. E.g. annotating thousands of gene functions or trait descriptions. Furthermore, new ontologies are regularly released, while older ones are updated by introducing new terms and obsoleting others, creating ongoing annotation tasks.

These challenges have led to the development of various auto-annotation tools. NCBO Bioportal provides the online *Annotator* tool [6] (formerly known as Open Biomedical Annotator), while EMBL-EBI provides a similar online tool known as *Zooma* [7]. Others include Ontobee[8], ontoFast [9] and Gene2Function [10]. Most of these tools use a form of text-mining, which limits their ability to find an appropriate term when the term label does not contain exact words contained in the query item, or when a concept is not succinctly encapsulated in the query item. The primary shortcoming of these tools, when compared to a human domain expert, is the lack of semantic understanding of the meaning of the query item and of the ontology term labels.

Large language models, such as GPT, are a recent innovation that move beyond text -mining and into the realm of semantic understanding, by modeling the relationship between words and phrases on a massive scale. LLMs show an increasing ability to find key concepts within complex writing [2], despite varying tense, tone, sentence structure and ordering of subjects. Therefore auto-annotation of scientific text descriptors with ontology terms is a task that plays to the strengths of LLMs. Here we describe a series of workflows incorporating OpenAI GPT’s [11]capabilities to annotate hundreds of *Arabidopsis thaliana* and forest tree phenotypic observations with ontology terms, aiming for results that resemble manually curated annotations. These workflows make use of an LLM to intelligently parse phenotypes into short concepts, followed by finding appropriate ontology terms via embedding vector similarity or via Retrieval-Augmented Generation (RAG) [12]. The RAG model is a state-of-the-art approach that augments conversational prompts to the LLM with context-specific data to empower it beyond its pre-trained parameter space. We evaluate the automated workflows against manually curated annotations using various performance metrics, including semantic similarity.

## Materials and Methods

The goal of this analysis was to improve the automated annotation of plant phenotype or trait descriptors with terms from the Plant Trait Ontology (TO) [13] by using an LLM. Starting from the existing auto-annotation approach that uses text-mining between a raw descriptor and ontology term labels, we tested three potential improvements: i) Using an LLM to split the raw descriptor into multiple concepts prior to annotation; ii) using LLM embedding-vector similarity for the annotation process; iii) finding candidate TO terms using (i) and (ii) above, then asking the LLM to select the best ones for annotation (RAG).

### Phenotype and Trait descriptors to be annotated

We aimed to auto-annotate descriptors from three plant phenotype sources. The first source is TAIR’s database [14] of phenotypes observed in mutant lines of the model plant *Arabidopsis thaliana*. A mutant line can produce one or more measurable phenotypic outcomes relative to wild-type. The TAIR database contains 19,235 phenotypic records for 7,551 genes (mutant lines), where each record may describe multiple trait observations. For example: *“AT1G34190 Shorter root length; reduced hypocotyl length after exposure of Antimycin”* is a single descriptor containing two observed phenotypes, both of which are part of a broader response to Antimycin (AA). The second source is the 1001 Genomes AraPheno database [15], which collates trait data measured on subsets of the 1001G Arabidopsis diversity population from hundreds of studies. Each trait has a short and often cryptic name, plus a longer description of how the trait was measured. The third source is the TreeGenes database [16] of traits aggregated from over 400 population-level studies in over 460 tree species. Each study investigated one or more traits, usually in the context of GWAS or adaptation, and a subset of those studies provide metadata about their measured traits.

### Gold-standard annotations

The TAIR mutant phenotypes were not already annotated with TO terms, so we manually annotated 100 randomly selected phenotypes to create a gold set of TO annotations for testing and evaluation. The Plant Trait Ontology includes some terms belonging to other ontologies (e.g. several PO, GO and CHEBI terms), so we avoided these and ensured that all gold annotations were terms with “TO” curies. Similarly, the TreeGenes traits were not already annotated with TO terms so we manually annotated the trait descriptions provided in study metadata files to create a gold set. We discarded any climatic and geographic traits, plus metabolite traits produced by mass-spectrometry as most of these are not describable by Plant Trait Ontology. Finally, the AraPheno traits were already annotated with a single TO term each, which we used for the gold annotation while acknowledging that a single term was often inadequate in capturing the full implications of the trait descriptor.

### Auto-annotation workflows

We evaluated the use of both text-matching and LLM embedding-vector similarity for auto-annotation of verbose descriptors. Additionally, we tested these annotators with an intermediate step where an LLM parsed the initial descriptor into multiple short concepts prior to the auto-annotation step (see Figure 1). Finally, we combined the outputs of the embedding workflows with the natural language capabilities of the LLM in a RAG approach. The five workflows in Figure 1 are explained in detail below.

**Fig 1.**
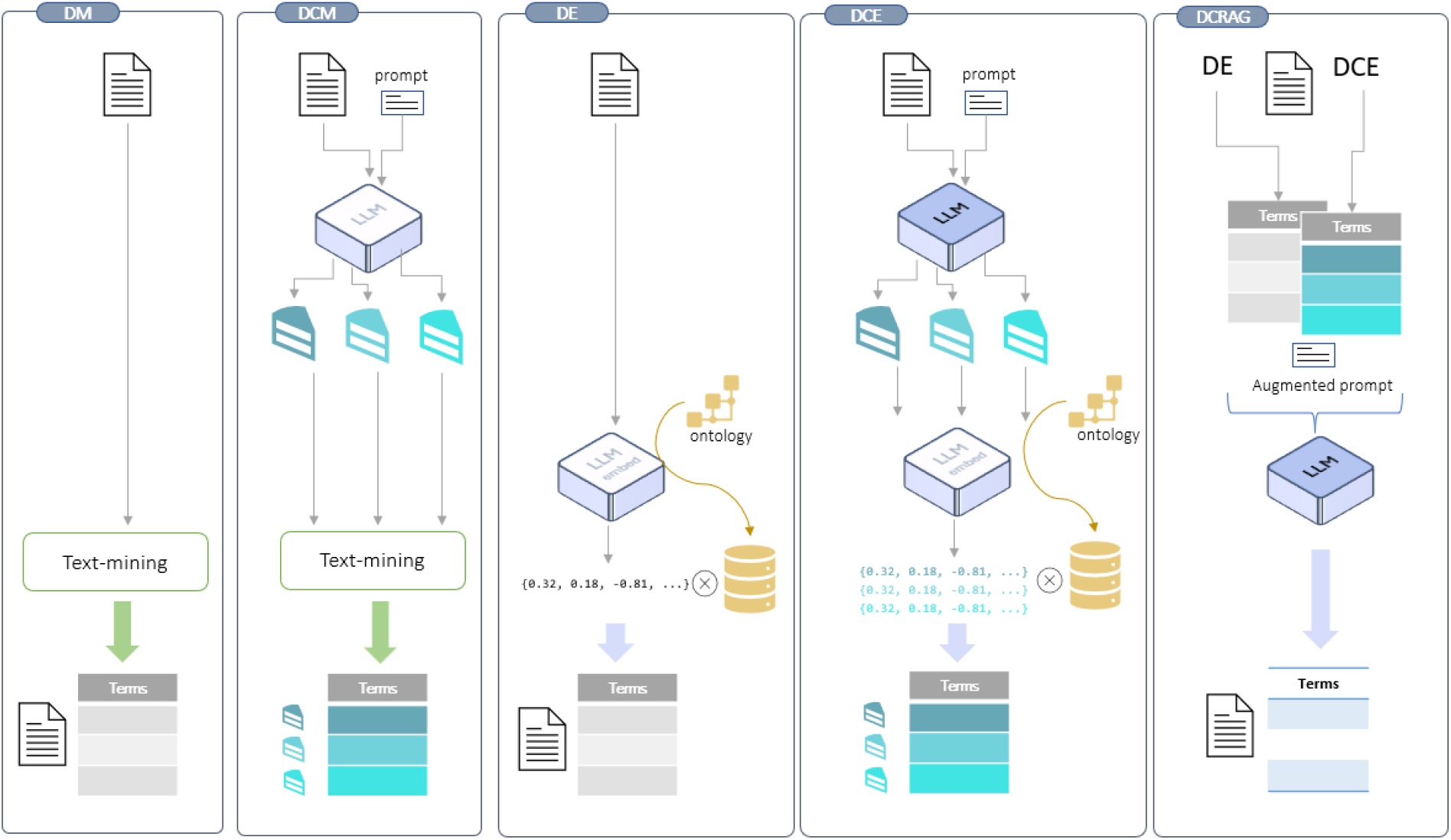
five auto-annotation workflows. Each workflow begins with a descriptor text input. i) DM sends the descriptor to a text-mining tool which annotates it with a set of ontology terms. ii) DCM asks the LLM to preprocess the descriptor into multiple short concepts (blue pieces), which are then annotated by the text - mining tool. iii) DE uses the LLM to get an embedding vector of the descriptor, which it then compares to a pre-calculated database containing embedding vectors of all ontology terms. The descriptor is annotated with the terms with the highest vector similarity to the descriptor. iv) DCE performs the same LLM preprocessing step in DCM to obtain concepts. DCE then uses the LLM to get an embedding vector of each concept, which it then compares to the embedding vector database of all ontology terms. The descriptor is annotated with the terms with the highest vector similarity to one or more concepts. v) DCRAG first runs the DE and DCE workflows to get a list of candidate ontology terms. It then asks the LLM to choose the most appropriate terms for the descriptor from the list of candidates.

### Descriptor to Text-Mining (DM)

This is the baseline method for auto-annotation. Here we provided each raw text descriptor as input to the online Zooma tool as a representative of text-mining algorithms, and requested annotation with the Plant Trait Ontology (TO). We removed any terms provided by Zooma that were not explicitly of the “TO:” curie that we used for the gold annotations for that dataset.

### Descriptor to Concepts to Text-Mining (DCM)

Most mutant line descriptors from TAIR have long, verbose descriptors, while AraPheno and TreeGenes traits are typically shorter but still can be verbose. Often many phenotypic observations or traits are described in the one descriptor. Here, rather than trying to annotate an entire raw descriptor in one go, we used an R wrapper of the OpenAI API [17] to ask the GPT-4o LLM to parse the descriptor into its major concepts, plus to provide three alternative short phrases for each of those concepts. The GPT prompts used for this task, which were slightly customized for each dataset, are available in **Supplementary S10**.

For each resulting concept we then concatenated the concept’s text and its three alternative phrases into a single comma-delimited text string, and then provided the string as input to Zooma for text-mining auto-annotation. Thus, if a raw descriptor could be broken into three separate concepts by GPT, Zooma would be used three times for that descriptor. A real example is given below.

#### Descriptor

“ABA hypersensitivity of guard cell anion-channel activation and stomal closing”

GPT detects three concepts (underlined here) with three alternate phrases each:

“<u>ABA hypersensitivity</u>, abscisic acid response, plant hormone sensitivity, guard cell regulation”

“<u>guard cell anion-channel activation</u>, anion transport, guard cell function, stomatal movement”

“<u>stomal closing</u>, stomatal closure, leaf gas exchange, plant water regulation”

The three strings above are used as three inputs to Zooma for the original descriptor.

### Descriptor to Embedding (DE)

An embedding is a numerical representation of a text input (or other data) that encodes the parameter space of the model for that input. It is a vastly compressed representation compared to the massive parameter space involved in large language models (e.g. a numeric vector of length 1024 rather than billions). Since embedding vectors encode a high-dimensional location in the language model’s parameter space, similar embeddings typically convey similar semantic concepts. For example “the quick red fox” and “a fast scarlet vixen” share no words in common but semantically describe highly similar concepts. Their GPT embedding vectors have a cosine similarity of 0.66. In contrast, “the quick red fox” and “four loaves of bread” have embedding similarity of only 0.21.

OpenAI provides a function to request embedding vectors for any input text. We first used this to get an embedding vector for each of the ontology terms (1671 TO terms), where the input for an ontology term was the concatenation of its text label and description. This formed our embedding vector database. Next we obtained an embedding vector for the raw descriptor, calculated its cosine similarity to each ontology term in the vector database, and selected the top four ranked TO terms with cosine similarity > 0.40 as annotations. All embeddings were calculated using the “text-embedding-3-large” model, which was the most advanced OpenAI embedding model available at the time of the study.

### Descriptor to Concepts to Embedding (DCE)

While the DE approach used embeddings of the raw verbose descriptors, in this approach we instead obtained embeddings of the GPT concepts described in the DCM workflow above. So, for each descriptor we obtained an embedding vector for each of its major concepts determined by GPT-4o. We calculated cosine similarity between the concepts’ embedding vectors and the ontology terms’ embedding vectors in the same way as the DE workflow, and then selected the single most similar ontology term for each concept (where cosine similarity > 0.40). Since long descriptors can be parsed into multiple different concepts by GPT-4o, this workflow could potentially auto-annotate a descriptor with many TO terms.

### Descriptor to Concepts to Retrieval Augmented Generation (DCRAG)

Embeddings are powerful tools but do not make use of LLMs natural language processing capabilities. However, if GPT-4o is asked directly to annotate a phenotype with ontology terms using pre-trained knowledge it tends to generate false term IDs and descriptions, so it is not trustworthy as an auto-annotator. The LLM needs to be provided with the dictionary of terms and definitions in the context window so it does not stray from the task. This is the domain of Retrieval Augmented Generation (RAG).

RAG is an established way to extend and focus the capabilities of an LLM. If an application needs the LLM to be aware of custom information then the LLM must be provided with that information as context in the conversation. Often the custom information is too large to be wholly included within the limited context space (such is the case with the Plant Trait Ontology which has over 1600 terms and their descriptions), so only the most relevant parts are retrieved from a local repository and then included in the conversation with the LLM.

In the DCRAG workflow we used the embedding similarity approach from the DE and DCE workflows to form a list of candidate TO terms. These are terms that, according to their embedding vector similarity with the descriptor or its concepts, have the best chance of being good annotations for the descriptor. For DCRAG a cosine similarity >= 0.35 (rather than 0.40) is used to grab a greater amount of TO terms. The list of candidate TO terms is concatenated onto a new LLM conversation prompt that asks the LLM to act as a plant biologist and choose the best terms from the candidates (**see Supplementary S10)**. The motivation behind this approach is to use the LLM’s demonstrated reasoning capabilities to intelligently winnow down the candidate TO terms, hopefully removing those that have a high embedding similarity with the descriptor but are actually false positives given the full context.

### Evaluating Workflows

In each workflow we compared the set of auto-annotated terms to the set of gold standard terms for each descriptor using Jaccard similarity and semantic similarity (semsim). Jaccard provides a score between 0 (no terms in common between the sets) and 1 (a perfect set of terms that is identical to the gold standard). However, given that ontologies are structured in a hierarchical graph, there are often terms existing at higher or lower levels in the graph that could be considered as valid annotations. The inability to positively score matches between similar (but not identical) terms is a shortcoming of Jaccard, and of any other similarity measure based on intersection of term IDs. Conceptually related terms with different IDs have a jaccard similarity of zero.

Semantic similarity, on the other hand, provides a more nuanced score between 0 and 1 that considers the structure of the ontology graph. Two sets of ontology terms can have no IDs in common yet still obtain a high similarity score if their terms are strongly connected in the graph. We used the *ontologySimilarity v2*.*7* [18] package in R to calculate Lin semantic similarity scores between sets of terms. Additionally, we calculated the precision and recall of auto-annotated term sets with respect to the gold term sets. All analysis was performed using R v4.3.0.

### Semantic similarity with random TO terms

It is difficult to know how easy or hard it is to attain a certain semantic similarity score, especially as the score can be affected by the size of the term sets. For example, is semsim = 0.62 a strong score when calculated between a set of 3 gold terms and a set of 4 auto-annotated terms? To answer this we took each descriptor and generated 100 sets of random TO terms of equal size to the set that was auto-annotated by a given workflow. We were thus able to calculate a ‘null’ distribution of semsim scores for each descriptor to compare the expected semsim of an unskilled auto-annotator to the semsim from the AI workflows.

## Results

### Gold standard annotation of descriptors

We manually annotated 100 randomly selected phenotypes from the TAIR mutant line dataset. On average, each phenotype received 2.14 TO terms, with the maximum being 8 terms. Four phenotypes could not be manually annotated with any TO terms so they were assigned *NA*. For the TreeGenes dataset, 224 traits were manually annotated in a similar manner to the TAIR phenotypes. On average the traits were annotated with 1.26 terms, with a maximum of 4. For the AraPheno dataset, the 231 unique trait descriptors were already annotated with a single TO term each. The complete gold annotations can be viewed in **Supplementary Tables S1-S3**.

### Parsing concepts with GPT

We asked GPT-4o to parse each input descriptor into concepts and provide three alternative short phrases for each concept. GPT appeared to be highly adept at this task. As an example, the TAIR descriptor “ABA hypersensitivity of guard cell anion-channel activation and stomal closing” was parsed into three concepts, each with three alternate phrases:

1. ABA hypersensitivity: abscisic acid response, plant hormone sensitivity, guard cell regulation
2. guard cell anion-channel activation: anion transport, guard cell function, stomatal movement
3. stomal closing: stomatal closure, leaf gas exchange, plant water regulation

The descriptors varied in length within and between the datasets, and there was a strong positive correlation between the length (word count) of a descriptor and the number of concepts determined by GPT (pearson’s *r*_TAIR_ = 0.71; *r*_TreeGenes_=0.72; *r*_AraPheno_ = 0.49). On average the TAIR descriptors produced the most concepts (3.44), followed by AraPheno (2.61) and TreeGenes (1.50). The entire set of concepts and phrases for all three gold datasets can be found in supplementary materials. Note that a small percentage of responses by GPT were badly formatted and needed to be remedied manually. All concepts generated by the LLM can be viewed in **Supplementary Tables S4-S6**.

### Auto-annotation

We tested five auto-annotation workflows (**Fig 1 and methods**) on each of the three gold-labeled datasets, and evaluated their performance using jaccard, precision, recall, and semantic similarity (semsim) metrics (**Table 1**).

**Table 1.** mean performance metrics for the five workflows in three datasets.

| dataset | workflow | preprocess | annotator | recall | precision | jaccard | semsim | r1 |
| --- | --- | --- | --- | --- | --- | --- | --- | --- |
| AraPheno | DCRAG | concepts | RAG | 0.372 | 0.269 | 0.269 | 0.598 | 0.372 |
|  | DCE | concepts | embedding | 0.268 | 0.177 | 0.177 | 0.511 | 0.268 |
|  | DCM | concepts | text-mining | 0.199 | 0.113 | 0.113 | 0.479 | 0.199 |
|  | DE | none | embedding | 0.355 | 0.089 | 0.089 | 0.575 | 0.355 |
|  | DM | none | text-mining | 0.091 | 0.071 | 0.071 | 0.270 | 0.091 |
| TAIR | DCRAG | concepts | RAG | 0.575 | 0.480 | 0.417 | 0.769 | 0.370 |
|  | DCE | concepts | embedding | 0.475 | 0.397 | 0.340 | 0.743 | 0.310 |
|  | DCM | concepts | text-mining | 0.411 | 0.332 | 0.279 | 0.627 | 0.250 |
|  | DE | none | embedding | 0.314 | 0.143 | 0.121 | 0.596 | 0.200 |
|  | DM | none | text-mining | 0.134 | 0.150 | 0.111 | 0.242 | 0.100 |
| TreeGenes | DCRAG | concepts | RAG | 0.583 | 0.517 | 0.494 | 0.731 | 0.558 |
|  | DCE | concepts | embedding | 0.388 | 0.362 | 0.342 | 0.664 | 0.366 |
|  | DCM | concepts | text-mining | 0.480 | 0.335 | 0.320 | 0.628 | 0.451 |
|  | DE | none | embedding | 0.517 | 0.146 | 0.144 | 0.663 | 0.496 |
|  | DM | none | text-mining | 0.228 | 0.163 | 0.158 | 0.399 | 0.219 |

The base approach of providing the raw descriptor to a text-mining annotation tool (DM) gave poor results in all datasets with all metrics. DM was generally unable to auto-annotate the exact terms found in the gold annotation, as evidenced by low mean recall values across each dataset (TAIR = 0.134; TreeGenes = 0.228; AraPheno = 0.091) and the fact that only a small proportion of descriptors had their entire set of gold terms recalled in full (r1: TAIR = 10%; TreeGenes = 21.9%; AraPheno= 9.1%). There was no single evaluation metric in any dataset where the DM approach was the best, and it was the worst performer in 12/15 such evaluations (**Supplementary table S8**).

The relatively poor performance of DM could be attributed to either its use of raw descriptors as input, or to its use of text-mining for auto-annotation. These aspects were addressed with the DCM workflow (which replaces the raw descriptor input with LLM-parsed concepts), the DE workflow (which replaces text-mining with LLM embedding-vector similarity), and the DCE workflow which does both. The results show that it is beneficial to use concepts as input instead of the full descriptor, or embedding-vector similarity as the auto-annotator. Using both together offers further improvement still (**Fig 2a**). The DCRAG workflow, however, was clearly the best as it scored the highest for every metric in every dataset (**Table 1**). Notably, DCRAG achieved much higher precision than all others while also achieving the highest recall, validating the use of an LLM’s natural language processing capabilities to accurately refine down a list of candidate ontology terms. Furthermore, DCRAG was often able to retain the exact same set of annotations as the manually curated gold annotations (**Fig 2b**), achieving this feat for 39% of the TreeGenes descriptors.

**Fig 2.**
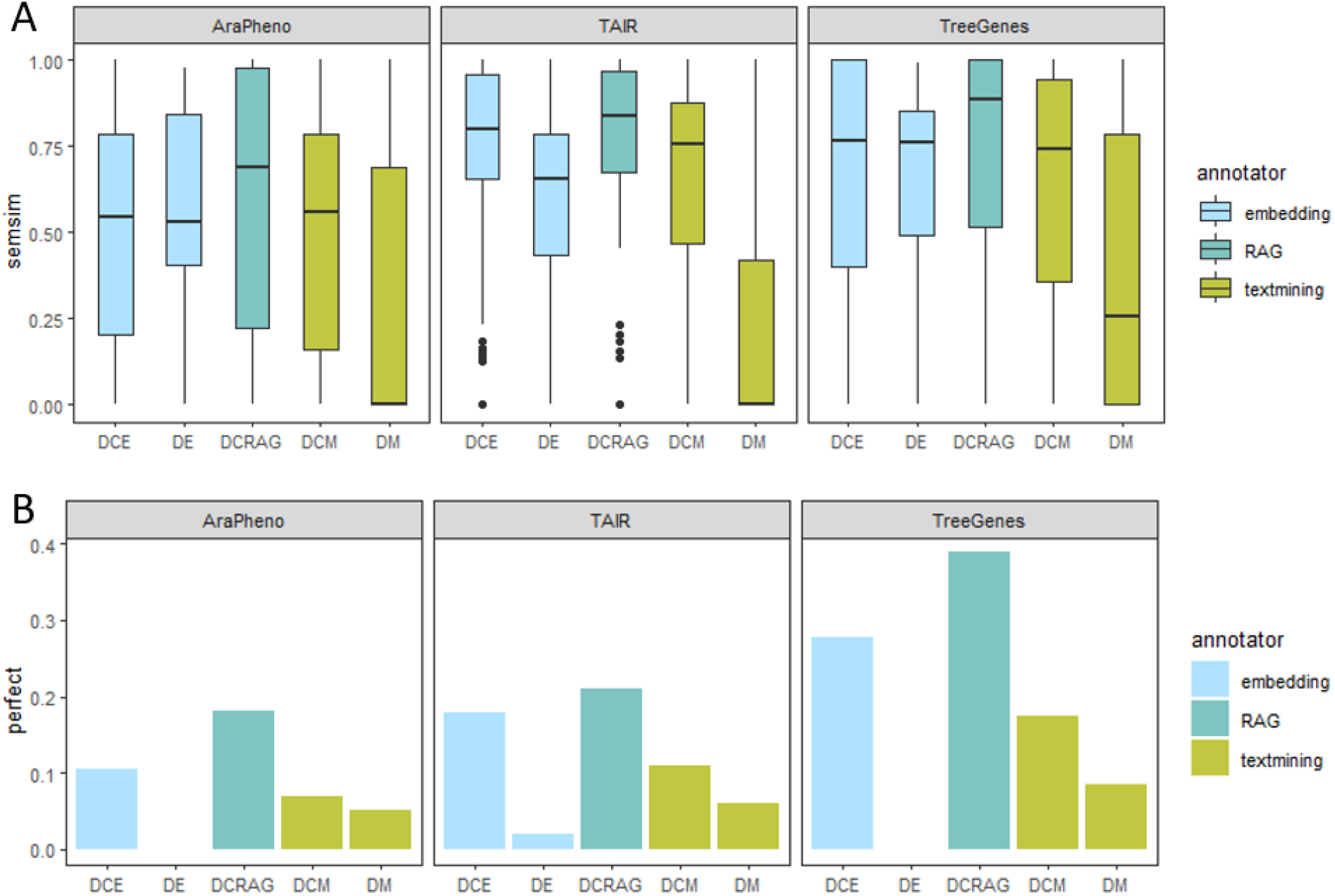
performance of workflows. A) Each boxplot represents the distribution of semsim scores from all the descriptors in the given dataset. The semsim score is a measure of how similar the auto-annotations are to the gold annotations for a descriptor, without relying on exactly matching ontology term IDs like Jaccard. B) The proportion of descriptors for which a workflow made perfect annotations (i.e. Jaccard = 1).

To understand how DCRAG improved auto-annotation performance we plotted the joint distribution of semsim scores from DCRAG and each competing workflow. In Figure 3, brighter regions of green-yellow indicate a high density of descriptors with similar score outcomes. If the brighter regions are above the diagonal line then the workflow on the y-axis tends to improve the performance for those descriptors compared to the workflow on the x-axis, while below the line indicates the opposite. For example, in each dataset the DM workflow produces a high-density region of poor semsim scores (0 - 0.25) which generally indicates a large number of descriptors with incorrect annotation. The DCRAG workflow, however, majorly improves the annotations for many of these descriptors, indicated by the green and yellow regions in the upper left of the panels in the top row. In the AraPheno dataset there are a number of low-scoring descriptors that could only be marginally improved by DCRAG. DCRAG also has an impact on descriptors that already scored reasonably well with the other workflows, as shown by brighter regions in the middle and top right of many panels that appear above the diagonal line. These are particularly noticeable against the DE and DCM workflows, but less so against DCE which performs strongly in the semsim metric and is hard to improve upon. DCRAG, notably, dominates DCE in both recall and precision, so its subtle improvements in semantic similarity score are generated by the selection of fewer and more accurate TO terms, which is highly desirable for auto-annotation.

**Fig 3.**
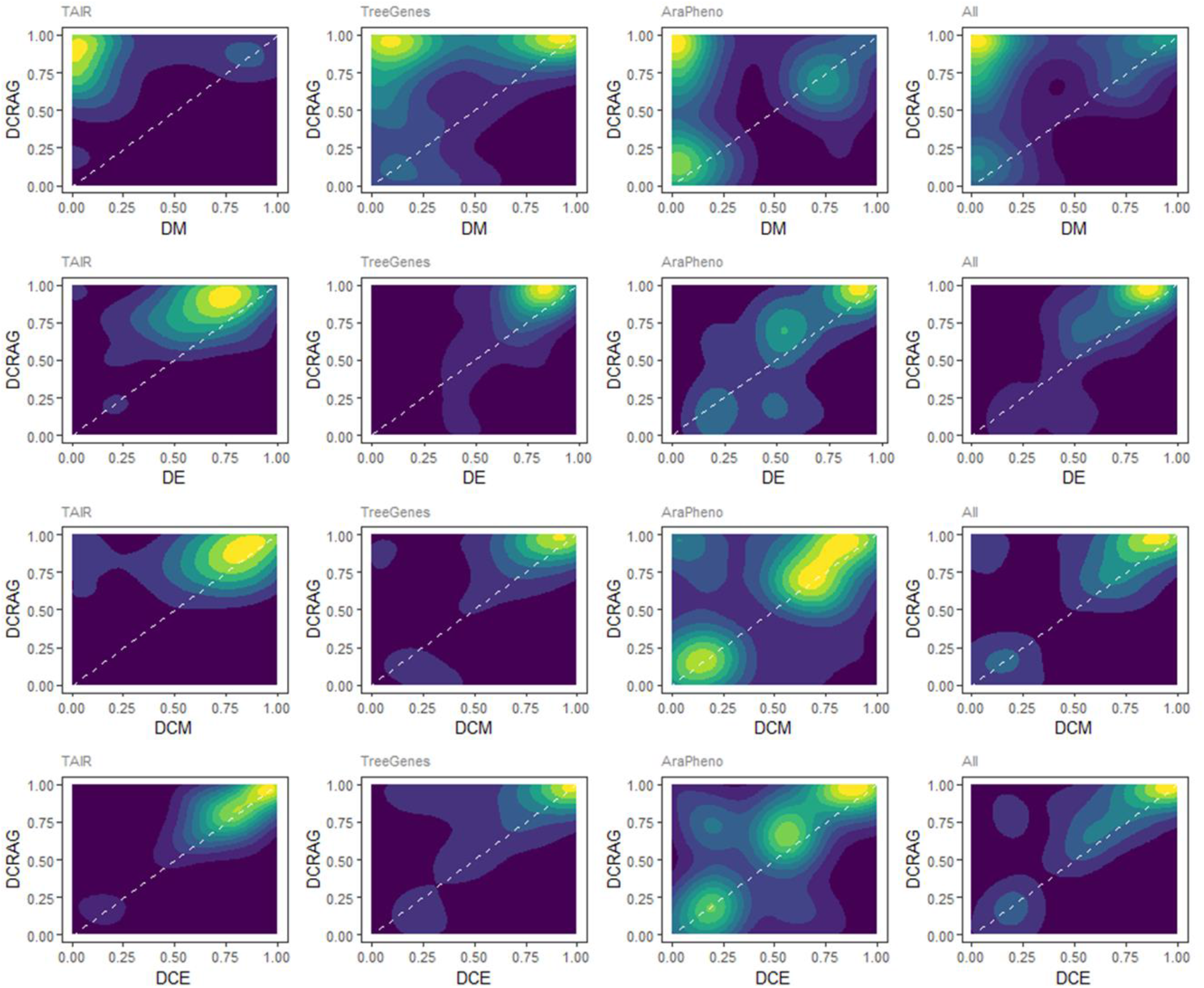
comparing semsim performance of DCRAG to other workflows. Each panel shows the joint distribution of semsim score for descriptors annotated by two competing workflows. Brighter regions of green-yellow indicate a high density of descriptors with similar score outcomes. The diagonal line represents identical performance, so bright regions above the line indicates the workflow on the y-axis performs better than the workflow on the x-axis for a dense group of descriptors.

### Semantic similarity with random TO terms

We calculated a ‘null’ distribution of semsim scores for each descriptor by randomly selecting TO terms for each descriptor 100 times and scoring the similarity to the gold terms. This let us compare the semsim of an unskilled auto-annotator to the semsim from the AI workflows (**Figure 4, Supplementary Table S11**).

**Fig 4.**
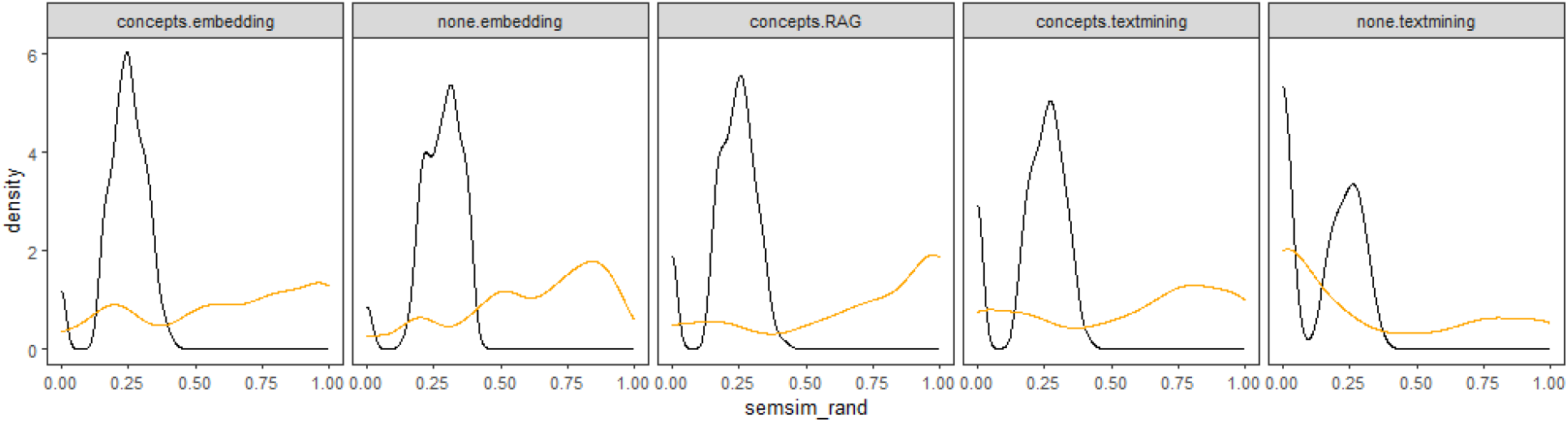
workflow performance versus random (unskilled) annotator. In each panel the black line shows the distribution of semsim scores when TO terms were selected at random, while the orange line shows the distribution of scores by the workflow. The number of terms selected at random for a descriptor was always the same as the number of terms annotated by the workflow. This was repeated 10000 times, so the black line is a mean distribution.

The random (unskilled) auto-annotator achieved a mean semsim of 0.22 and a maximum of 0.42. Semsim scores were correlated with the number of terms auto-annotated to a descriptor, but this effect was much stronger for random sets than for the AI workflows (random: *r* = 0.66; workflows: *r* = 0.31). This indicates that it is easier to get a higher semsim score by making more annotation “guesses”, but the performance of the AI annotators is not as dependent on this effect as the random annotator is.

Since semsim score is somewhat confounded by the number of terms in the annotation set, and each workflow varied in the number of terms annotated to each descriptor, we used fold-change between a workflow’s semsim score and its random semsim score as a robust indicator of skill. DCRAG was once again the best performer in all datasets, with a semsim score up to 3.4 fold greater than random annotation for the TreeGenes dataset (**Supplementary Table S11**).

### Annotatingthe un-annotatable

The TAIR dataset contained four descriptors without gold annotations as there were no suitable TO terms for them. It is a challenge for a workflow to refrain from assigning any terms to these descriptors. The DM workflow correctly assigned zero TO terms to 3 out of the 4 un-annotatable descriptors, but that must be viewed within the context that DM also assigned zero terms to 55 other descriptors out of the 100 in the TAIR dataset. In comparison, DCRAG correctly assigned zero TO terms to 3 out of the 4 while assigning zero terms to just two other TAIR descriptors. DCE was the best performer here as it assigned zero terms for all 4 un-annotatable descriptors plus just one other TAIR descriptor (**Supplementary Table S12**). This indicates that AI workflows like DCRAG and DCE are capable of recognising and annotating plant traits within broader text while also recognising when no meaningful trait is present.

## Discussion

Annotating scientific text with ontology terms is a difficult task for humans with domain expertise, let alone for automated tools. We demonstrated this by using a text-mining tool to auto-annotate hundreds of plant phenotypes and gene functions. For these inputs, which were not written with auto-annotation in mind, the resulting annotations rarely included the gold (curated) terms, even for the simplest, shortest, descriptors found in the TreeGenes traits dataset. This is because text-mining approaches can fail when words do not match exactly. For example, Arabidopsis trait #278 (https://arapheno.1001genomes.org/phenotype/278/) found in the AraPheno database is described as “iron concentrations in leaves”. NCBO’s Bioportal Annotator finds no matching TO terms, even when using the “match partial words” option. Similarly, Zooma was unable to match “iron concentrations in leaves” to any TO terms, instead focusing on the word “leaves” and annotating it to Plant Ontology (PO) terms for leaves. Yet if we slightly modify the descriptor by removing the ‘s’ in “concentrations” then both tools find the correct match to TO:0006049 “iron concentration”. While some text-mining algorithms may incorporate more sophisticated rules that can handle this situation, an LLM would not be so easily tripped up by plurals, tenses or word ordering, nor get “distracted” from the subject of the sentence.

### Concept parsing offers easy improvements

It is tempting to use an LLM like GPT-4o to directly annotate phenotypes with TO terms by simply prompting it to do so. However, when we tried this, GPT-4o intermittently “hallucinated” TO terms to match the concepts it intrinsically found in the phenotype text. Regardless, it was clear that the GPT-4o model was highly capable of finding the key concepts in the phenotype despite the complex and often abstract language being used. Additionally, by asking GPT to rephrase a concept multiple times, the annotation of that concept could be approached from different angles which gives downstream tools a greater chance of finding the correct ontology terms. This was evident from result comparisons between workflows. DCM and DCE included a concepts-parsing preprocess step prior to annotation and performed consistently better than their DM and DE counterparts.

### Retrieval Augmented Generation is a powerful tool for annotation

The DCRAG workflow can be viewed as a logical extension of the DE and DCE workflows. While DE and DCE are a major improvement over DM, their final selection of ontology terms relies on an arbitrary cutoff of embedding vector similarity (e.g. cosine > 0.40). Make the cutoff too low and too many false positive terms will result; make the cutoff too high and many true positive terms will be left out. It is telling, for example, that the DE workflow achieved a strong average recall of 0.517 for the TreeGenes dataset but average precision of only 0.146. Clearly, for many TreeGenes descriptors, a great deal of true positive ontology terms made it through the cosine similarity cutoff but they were accompanied by even more false positives. A subject expert would be able to review this list of terms and discard those that don’t fit, improving precision without sacrificing recall. In the case of DCRAG the LLM was instructed to act as that expert, and it did so with a proficiency that resulted in considerably higher scores in every metric and dataset. DCRAG was able to select perfect annotations (i.e. Jaccard = 1) for 18%, 21% and 39% of all descriptors in the AraPheno, TAIR and TreeGenes datasets respectively. While this is not enough to confidently replace a human annotator, it does suggest that LLM-based workflows may have the potential to do so in future.

A limitation of DCRAG is the number of candidate ontology terms to include as context in the augmented prompt. If the list gets too large GPT-4o may struggle to use their information accurately. We combined TO terms generated from the DE and DCE approaches using a cutoff of cosine > 0.35, which typically produced between 10 and 30 candidates. Using both DE and DCE outputs meant that some terms were candidates due to their similarity with the entire descriptor (the DE approach) while others were candidates due to similarity with one or more short concepts (the DCE approach). Thus, the LLM was given local and global options to choose from.

## Conclusions

Here we have shown that LLM-based approaches for auto-annotation of plant phenotypes offer an improvement over a text-mining approach. Concept parsing and embedding with an LLM helps to avoid the pitfalls of text-mining, can be automated using an API, and results in much improved auto-annotations. Further improvements are gained using a RAG approach to guide the LLM in finding the best ontology terms for annotation. In essence, our RAG workflow emulates that of a human annotator who would read the input text, understand the concepts within, peruse the target ontology for candidate terms, and then decide on which candidate terms are most appropriate based on the broader meaning and the concepts within the input text.

The accuracy of RAG annotations is by no means yet at a level to replace human experts, but can provide a high-throughput automated way to produce a first pass set of annotations for rapid curation by experts.

## Supporting information

Supp 1 - gold descriptors

Supp 2- gold concepts

Supp 3 - annotation results

## Key Points

- Large Language Models are well suited to the task of auto-annotating phenotype descriptions with trait ontology terms
- Using an LLM to deconstruct longer phenotypes into shorter concepts improves annotation accuracy
- A Retrieval Augmented Generation (RAG) model boosts the AI capability for annotation considerably

## Acknowledgements

Thanks go to Dr. Stephanie Conway for providing expert plant biology assistance when curating the gold annotations for the TAIR dataset.

## Funding

This work was funded by The Australian Research Council Centre of Excellence for Plant Success in Nature and Agriculture (CE200100015).

### Data Availability

All datasets used in this study are available as supplementary files. This includes phenotype descriptors, gold standard annotations, concepts parsed by LLMs, auto-annotations from each workflow and evaluation scores of the auto-annotations. Code to execute the DE, DCE and DCRAG workflows is available at https://github.com/dkainer/LLMannotator.

